# Imprecise counting of observations in averaging tasks predicts primacy and recency effects

**DOI:** 10.1101/2024.09.29.615676

**Authors:** Arthur Prat-Carrabin, Michael Woodford

## Abstract

Primacy and recency effects — wherein early and recent observations exert disproportionate influence on judgments — have long been noted in cognitive tasks involving the sequential presentation of information. In studies where human subjects make decisions based on the average of a sequence of numbers, recency effects are typically modeled phenomenologically through exponential discounting, while primacy effects are neglected altogether. Here, we exhibit the prevalence of both effects in such tasks, and propose that they result from the observer’s imprecision in their running tally of how many pieces of information they have received. If their approximate counting follows a central tendency — a typical Bayesian pattern — then past information is overweighted near the beginning of the sequence, while new numbers are overweighted towards the end of the sequence. Thus both primacy and recency effects are simultaneously predicted by this single mechanism. The model moreover nests exponential discounting as a special case in which the observer has no information about the count. The behavioral data suggests that subjects indeed misestimate the count of observations, with biases similar to those observed in numerosity-estimation tasks. Finally, we present evidence that the central tendency of subjects shifts towards lower counts in tasks with shorter sequence lengths, consistent with a Bayesian estimation of the counts. These findings provide new insights into the cognitive processes underlying serial-position effects in averaging tasks, with broader implications for other cognitive domains.

---

Information often comes one piece at a time. Individuals process and integrate this information, one piece after the other, to form judgments or make choices. In doing so, they frequently exhibit serial-position effects, and in particular primacy and recency effects, whereby early and recent information disproportionately influence their decisions. These effects have been observed for decades and across various modalities: e.g., in forming impressions of personality on the basis of a list of traits^1,2^, in memory recall^3–6^, causal judgments^7^, inference and prediction^8–11^, and perceptual decisions^12,13^. A kind of tasks in which each piece of information should clearly have the same influence on decisions is averaging tasks, in which subject are asked to make a judgment on the basis of the average of the magnitudes of stimuli presented to them sequentially. The magnitudes may be physical properties of the stimuli (e.g., length or orientation^14,15^), or numerosities, as in clouds of dots^16^, or presented as Arabic numerals^17–20^. Most of these number-averaging studies find recency effects and some find primacy effects. But the more recent studies, in any case, do not focus on serial-position effects, which are often treated as peripheral: primacy effects are not considered, and recency effects are typically captured in models through an exponential discounting of remote observations. This assumption has become so commonplace that it is rarely justified in detail, and the broader implications of serial-position effects are often ignored.

We aim to fill this gap by examining primacy and recency effects in number averaging tasks more closely. We exhibit the significance of these effects in a recently published dataset^20,21^, and we propose a single, plausible mechanism that accounts simultaneously for both effects. Simply put, we assume that when an observer receives the *n*^*th*^ number of a sequence, they are uncertain about how many numbers they have seen thus far (i.e., they are imprecise about *n*). Consequently, they are uncertain about how much weight this new number should carry in their updating of their running estimate of the average. If they estimate the count *n* with a central tendency (as is typical for Bayesian observers), then at first they will underestimate the weight of the new number (and thus overestimate the weight of the numbers already seen, resulting in primacy effects), but later in the sequence they will overestimate the weight of the new number (resulting in recency effects). Primacy effects and recency effects, by this account, thus indirectly reflect the central tendency of Bayesian observers. In the extreme case where the observer has no sense of the count of observations, our model reduces to the exponential-discount models commonly used in the literature. However, in more realistic scenarios where observers have a reasonable but imperfect estimate of the count, both primacy and recency effects naturally emerge.

The remainder of the paper is structured as follows. First, we analyze existing data from a sequential number averaging task to document the presence of both primacy and recency effects. Then we introduce our approach and demonstrate how it accounts for these effects. We then re-analyze the behavioral data under this new conceptual light and exhibit how subjects seem indeed to mis-estimate the count of observations, which in turn results in primacy and recency effects. We then predict that with shorter sequences, the crossover point of subjects’ central tendency should come earlier, and we find evidence for that in two other datasets. Finally, we conjecture in the Discussion how this approach could be applied to problems of memory recall. Together, our results suggest that serial-position effects find their origin in a suboptimal integration of new information that results from an imprecise estimation of the amount of prior evidence received.

## Results

### Primacy and recency effects in a sequential averaging task

We analyze the responses of human subjects in one of the averaging tasks documented in Ref. 20 (we are indebted to the authors of this study, who made their data available online^21^). In each trial of this task, the subject is presented with a sequence of *T* = 8 numbers randomly (and independently) sampled from a given distribution, and displayed for 350ms each. The subject is then asked to indicate whether the empirical average of these numbers is higher, or lower, than a reference value, *µ*, which is the mean of the distribution from which the numbers are sampled. The authors manipulate this distribution and study how this impacts the behavior of subjects; but here, we are interested in examining serial-position effects, i.e., whether the position of each number in each sequence modulates its influence on the decision of the subject, and thus we do not discuss the different experimental conditions. (In addition, the task we focus on is the ‘single-stream’ task run by the author; we briefly discuss further below the other, ‘dual-stream’ task that they implement.) More details on the task can be found in Ref. 20.

In order to estimate the influence of each number on subjects’ decisions, we fit a decision model in which an estimate of the average is compared, with noise, to the reference value, *µ*. Specifically, the model subject computes a weighted average of the presented numbers,

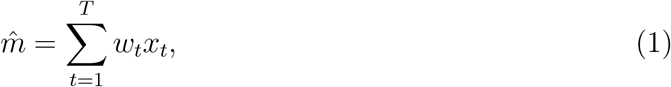

where *x*_*t*_ is the number presented in position *t*, and *w*_*t*_ is the weight applied to this number in the estimate of the average, 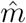. These weights sum to 1. The subject then compares this estimate to the reference value, *µ*, but with Gaussian noise whose standard deviation is *ν*. More precisely, the probability that the subject decides that the empirical average is greater than the reference value is 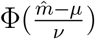, where Φ is the cumulative distribution function (CDF) of the standard normal distribution. We fit this model to the responses of each subject, by maximizing its likelihood.

A correct calculation of the average would apply to each number the same weight, *w*_*t*_ = 1*/T* = 0.125. We find instead that the weight of a number *x*_*t*_ in a subject’s estimate of the average strongly depends on its serial position, *t*. The weight applied to the first number (*t* = 1) is not significantly different from its correct value, but weights after that decrease with the position *t*, up to *t* = 4 where the median weight, 0.101, is significantly lower than the correct weight and lower than the weight applied by the subjects to the first number (Fig. 1). In other words, subjects exhibit a primacy effect. Later numbers, conversely, see their weights increase as a function of their position in the sequence, and the largest weight is applied to the very last number (median weight equal to 0.159, significantly greater than the correct weight, and greater than the weight applied by subjects to the number in fourth position; Fig. 1). In other words, subjects clearly exhibit both a primacy effect and a recency effect. Many studies model the recency effect through an exponential discount of more remote observations, and do not model the primacy effect (perhaps because it is smaller). We now present a model that accounts simultaneously for both.

**Fig. 1:**
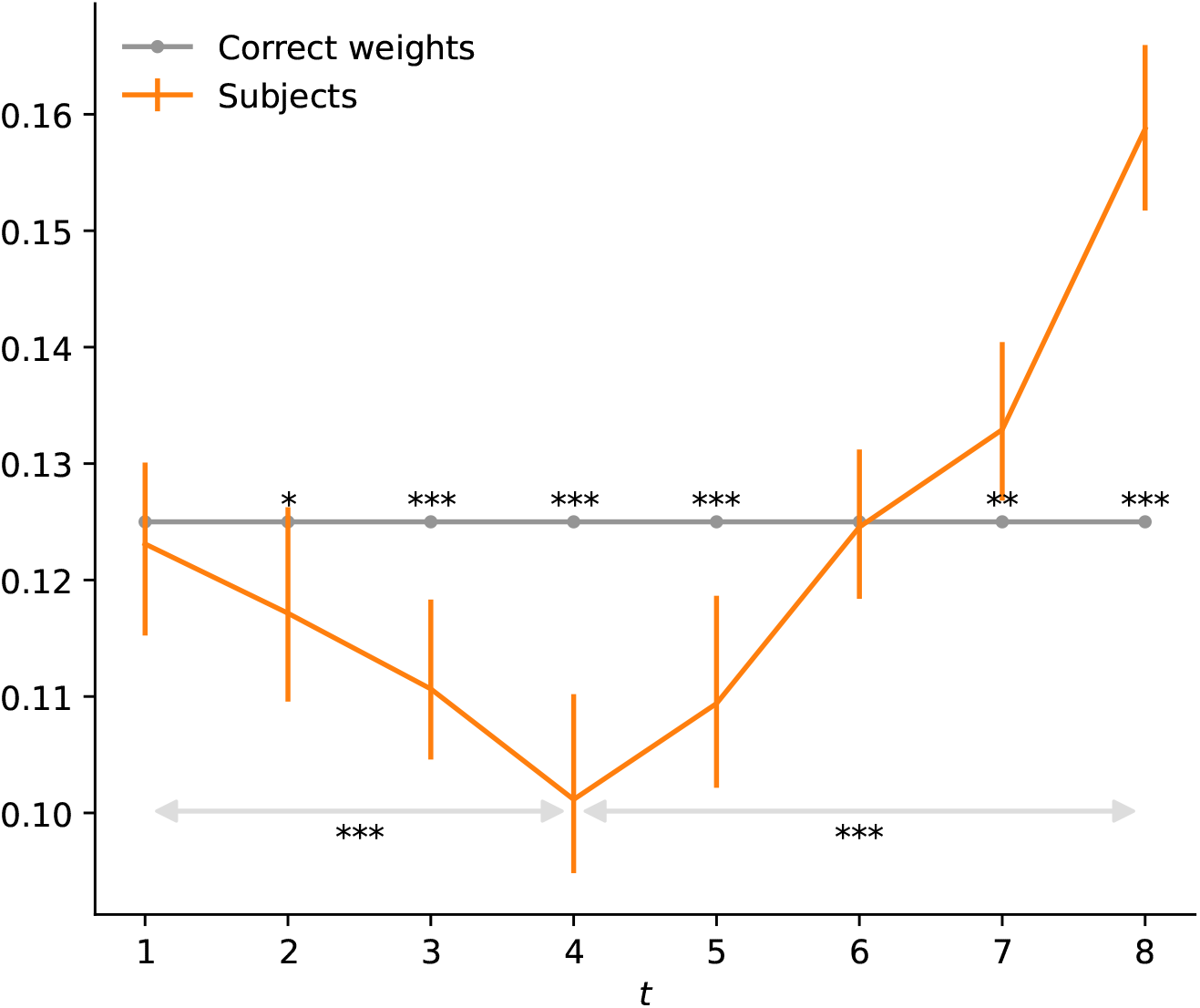
Subjects’ primacy and recency effects in an averaging task with sequentially presented numbers. Correct weight (grey line) and median of subjects’ weights (orange line) as a function of the position *t*. Stars indicate the p-values of permutation tests (30,000 permutations) of the equality between subjects’ weights and the correct weights (stars located above the correct-weights line), and between subjects’ weights at *t* = 4 vs. at *t* = 1 and *t* = 8 (*: *p <* .05, **: *p <* .01, ***: *p <* 0.001). Error bars indicate bootstrap-estimated 95% confidence interval.

### Sequential averaging with imprecise counts

We consider an observer who, like the subjects in the study presented above, is asked to indicate whether the empirical average of *T* numbers presented sequentially is higher, or lower, than the mean *µ* of the distribution from which these numbers are sampled. Because the observations of the numbers unfold in time, we assume that the observer computes the average in an ‘online’ fashion, i.e., by maintaining throughout the sequence an estimate of the average and updating it every time a new number is presented. We denote by 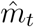 the estimate of the average after presentation of the number *x*_*t*_. We posit in addition that before seeing any number, the observer starts with an estimate equal to the mean of the distribution, i.e., 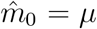 (this assumption will have however little consequence, as in most cases presented below this initial estimate will receive no weight in the final estimate of the average). We assume that the updated estimate is obtained as a weighted average of the preceding estimate and of the new number, as

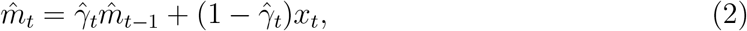

where 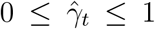. Such update rule indeed yields the correct average of the presented numbers if the weight applied to the preceding estimate is the correct one, 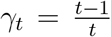. To compute the correct weight, one must thus remember the number of observations seen thus far, *t*.

Here we assume that the observer is uncertain about this count. As a result, they choose a weight 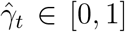 that may differ from the correct weight *γ*_*t*_. The observer may for instance receive a noisy signal, *r*_*t*_, about the observation number *t*, and choose for the weight 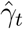 the expected value of the correct weight given the signal *r*_*t*_, i.e., the Bayesian posterior mean, 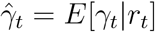. The behavior of such estimate depends on the specifics of the noisy signal and of the observer’s prior, but Bayesian estimates typically overestimate small magnitudes and underestimate large magnitudes (‘central tendency effect’). As the correct weight, 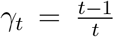, increases with the observation number *t*, the weight 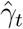 applied by the observer to the preceding average will typically exceed the correct weight at the beginning of the sequence, and fall below the correct weight at the end of the sequence. In other words, when near the start, past observations are overweighted; but near the end they are underweighted.

To investigate whether this results in primacy and recency effects, we need to examine the resulting weights of each number in the final estimate of the average: these result from the compounded effects of the weights 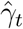 applied to the successive estimates of the average.

More precisely, the estimate of the average after the last observation is the weighted average

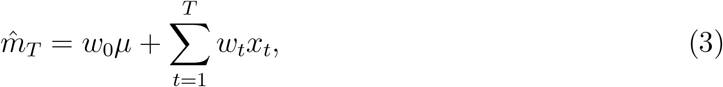

where the weight *w*_*t*_ of the number that appeared at time *t* ≥ 1 is

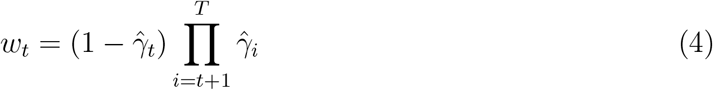

(with the convention that an empty product is equal to 1), and the weight *w*_0_ applied to the mean *µ* is

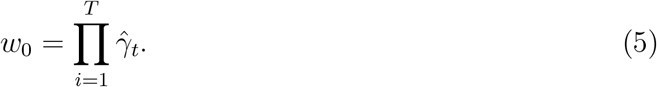

(Note that 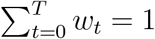.) We call ‘successive weights’ the weights 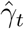 applied to the successive estimates of the average, and ‘number weights’ the resulting weights *w*_*t*_ on the numbers. With the correct successive weights 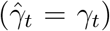, the correct number weights are obtained: *w*_0_ = 0, i.e., no weight on the mean *µ*, and 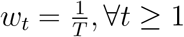, i.e., equal weights applied to each number in the estimate of the empirical average (Fig. 2, grey lines).

**Fig. 2:**
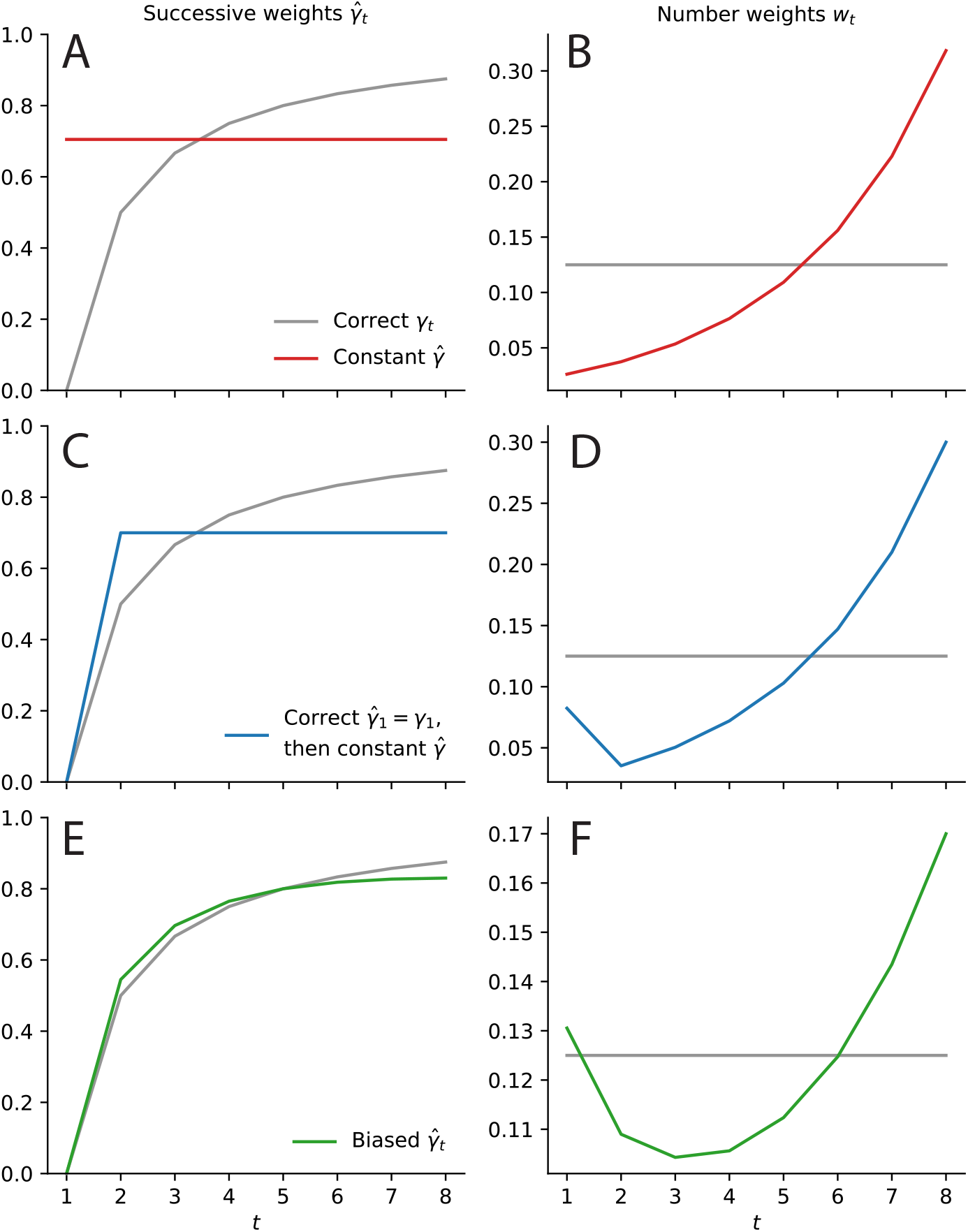
Imprecise estimates of the correct weight on past information yield recency and primacy effects. Successive weights 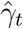 on the preceding average (left column) and the resulting number weights *w*_*t*_ (right column) as a function of number position, *t*, with constant successive weights (A, B, red lines), correct first successive weight 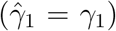 followed by constant successive weights (C, D, blue lines), biased successive weights (E, F, green lines), and correct weights (grey lines). The number weights *w*_*t*_ in the right column are obtained from the successive weights 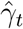 in the left column through Eq. 4.

As a first illustration, we note that the last observation receives the weight 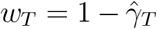. At the last observation, the weight applied to the previous average should be 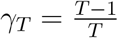, i.e., its highest possible value for a sequence of *T* numbers. But the observer, unsure about the correct value of this weight, relies on an estimate, which presumably is not greater than the maximal value. In most cases it will be lower, i.e., the observer chooses a weight 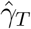 that underestimates the correct weight for the previous average, and thus a weight on the last observation, 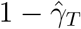, that overestimates the correct one. Therefore, the observer overweighs the last number in their estimation of the average, i.e., they exhibit a recency effect.

Beyond this simple illustration, the patterns of under- and overweighting of the numbers in the estimate of the average depends in non-trivial ways on how the weights applied to the successive averages, 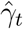, under- or over-estimate the correct weights, *γ*_*t*_. We thus consider a few illustrative choices of the successive weights 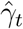, and examine the behavior of the resulting number weights, *w*_*t*_.

### No-information model

We start with the case of an observer who does not have access to any information that would allow them to estimate the current count of observations. Thus at every presentation of a new number, the observer is in the same state of knowledge (or lack thereof), and therefore they always choose the same weight. Thus 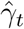 is a constant, which we denote by 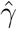 (Fig. 2A). The number weights are then

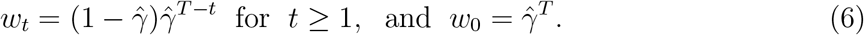

Thus the number weights *w*_*t*_, for *t* ≥ 1, amount to an exponential filter, by which more remote observations are more strongly discounted. Such exponential-discount model is commonplace in the literature on sequential averaging^16,19,20^. Here we have shown that an observer who seeks to update their estimate of an average, but who does not know how to weigh the incoming evidence relatively to preceding estimate, will exhibit exponential discounting. As each number is weighted less than the next 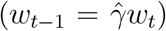, this observer exhibits a recency effect (Fig. 2B).

### No information on the count, except for the first

The observer just presented might seem an extreme case. In particular, even when presented with the very first number of the sequence, this observer is unable to recall how many numbers they have seen. We now turn to a different observer, one who knows upon seeing the first number that it is indeed the first. In other words, they correctly choose the initial successive weight as 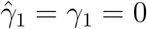(and henceforth we will consistently make this assumption; note that it implies *w*_0_ = 0). Starting from the second number, we assume once again that the observer does not remember how many numbers they have seen (except that they know it must be more than one, as otherwise they would know that it is one), and thus the successive weights they choose are here also a constant, 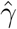 (Fig. 2C). This results in the number weights

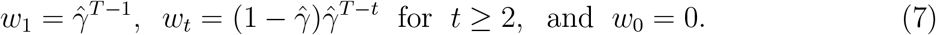

Here also the number weights yield an exponential discount, and thus a recency effect, but only for the numbers between *t* = 2 and *T*. As for the weight on the first number, *w*_1_, we note that 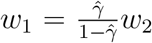, and thus whether the first number will be weighted more in the final average than the second number depends on whether 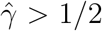. What weight 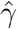 should our observer choose? The successive weights *γ*_*t*_ form an increasing function of *t*, and the observer knows that the count is not 1, and thus that it must be 2 or more: it would be reasonable, then, to choose a weight 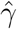 equal to or greater than *γ*_2_, the correct weight for the second number; and strictly greater than *γ*_2_, if *T >* 2. But *γ*_2_ is precisely equal to 1*/*2 (as it is the weight applied when taking the average of two numbers). Therefore, *w*_1_ *> w*_2_, i.e., this observer exhibits a primacy effect, in addition to the recency effect already mentioned (Fig. 2D). In short, we obtain both effects with an observer who knows that the initial number is indeed the first, but who is unable to recall the correct count after that.

### Biased estimates of the correct weights

The second observer just presented may seem more plausible than the first, but it may still seem improbable that as early as the second observation human subjects would entirely lose track of the count of observations. A possibility is that they receive a noisy signal *r*_*t*_ about the count *t* that allows them to derive a reasonably good, but not perfectly accurate, estimate of the correct count (except at the first observation, for which we assume that they accurately identify that it is indeed the first, i.e., 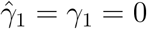, and thus *w*_0_ = 0). We assume that there is a value *τ* ∈ [2, *T*] such that for counts *t* greater than 2 but lower than *τ*, the observer on average overestimates the count *t*, while for counts above *τ*, on average the observer underestimates the count *t*. (The averages are taken over realizations of the random signal *r*_*t*_.) This ‘central tendency’ is a typical bias of Bayesian observers. Consequently, the successive weights 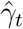 chosen by the observer are overestimations of the correct weights *γ*_*t*_ up to *τ*, and underestimations of the correct weight for *t > τ*. We call *τ* the ‘crossover point’. For the sake of simplicity, we assume that this bias is linear, as

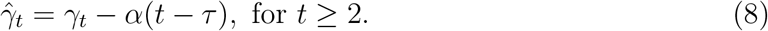

For the crossover point, we choose the middle value between 2 and *T* ; thus with *T* = 8 we choose *τ* = 5. At the crossover point, the chosen weight is correct, i.e.,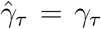. The successive weights chosen by this observer is shown in Figure 2E (for *α* = .015). As for the resulting number weights, they decrease from *t* = 1 to *t* = 3, after which they increase (Fig. 2F). In other words, this observer exhibits a primacy effect, followed by a recency effect; and the behavior of their number weights resemble that of the subjects (Fig. 1). We thus obtain a behavior similar to that of the subjects by positing an observer who correctly identifies that the initial observation is the first one, and whose estimation of the subsequent counts are imprecise, but consistent with Bayesian inference in that they overestimate smaller magnitudes and underestimate larger magnitudes.

### Subjects as imprecise counters

The analysis presented above suggests that the primacy effect and the recency effect found in subjects’ behavior do not result from two separate mechanisms, but stem instead from a single origin, the imprecision in subjects’ evaluation of the amount of evidence presented. This imprecision induces in turn the incorrect weight they apply to the new information, relatively to their previous estimate of the average. This mechanism prompts us to examine directly the successive weights 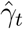 chosen by the subjects, instead of the number weights *w*_*t*_. (We note that while the weights *w*_*t*_ must obey the normalization constraint 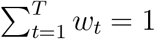, the successive weights 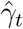 have no such constraint, and thus they cannot be identified without an additional assumption; in what follows we assume that 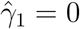, i.e., that upon seeing the first number the subjects know that the count is *t* = 1.) We find that the subjects’ successive weights are larger than the correct weights when the number *t* of observations is lower than 5, and smaller than the correct weights when *t* is greater than 5, with significant differences for *t* = 4, 6, 7, and 8 (Fig. 3, left column). This pattern of over- and under-estimation is thus similar to that of the model observer presented above, with a similar crossover point *τ* around 5.

**Fig. 3:**
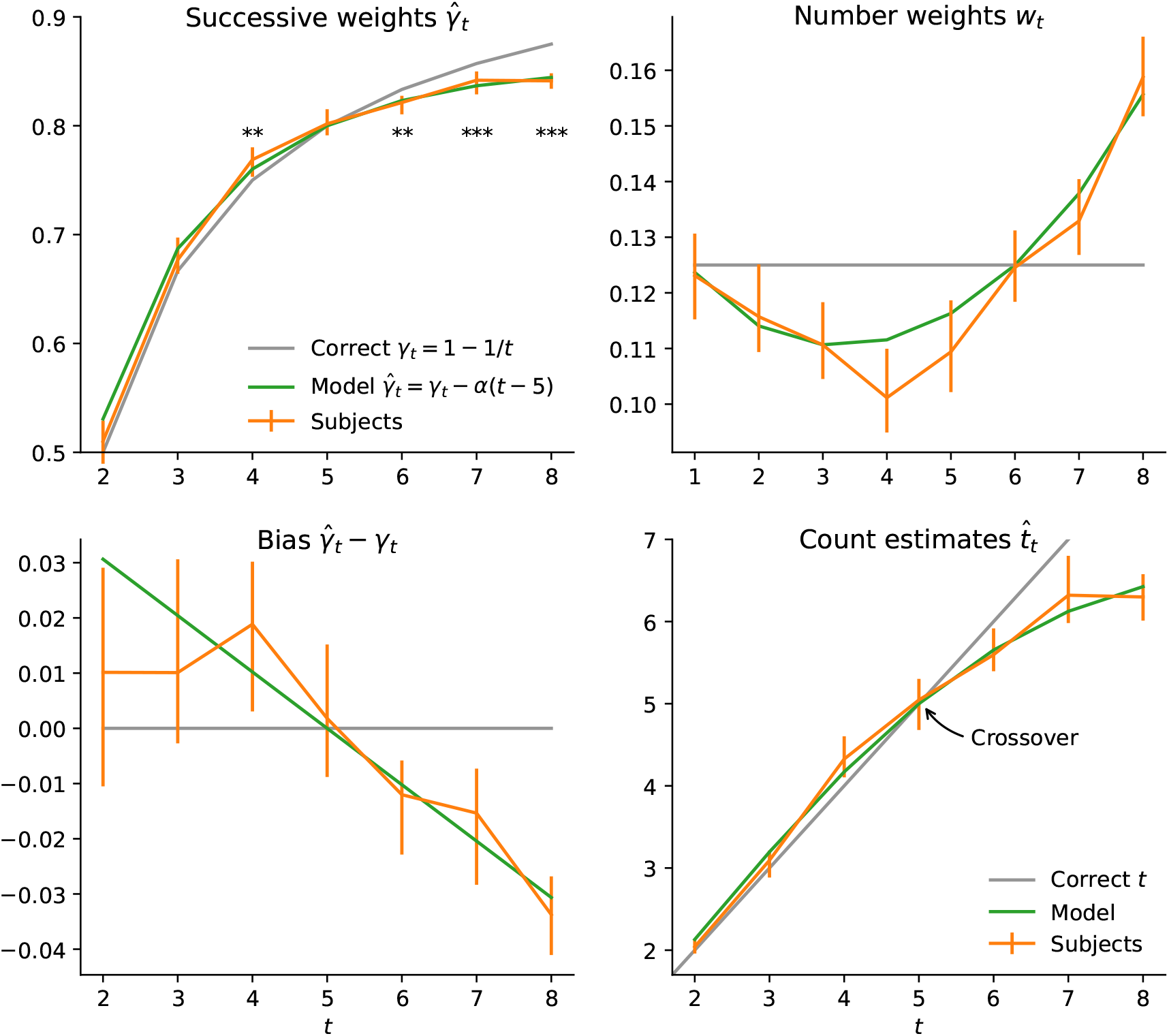
Subjects’ primacy and recency effects originate in their imprecise counts of seen observations. Correct (grey lines), biased model’s (green lines), and subjects’ median (orange lines) successive weights 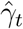 (top left), biases 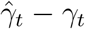 (bottom left), number weights *w*_*t*_ (top right), and approximate counts 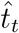 (bottom right), as a function of number position, *t*. Stars indicate the p-values of permutation tests (30,000 permutations) of the equality between subjects’ successive weights and the correct weights (**: *p <* .01, ***: *p <* 0.001). Error bars indicate bootstrap-estimated 95% confidence interval. The data in the top-right panel is the same as in Fig. 1.

We can also directly estimate the subjects’ approximate counts, 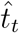, that yield their successive weights 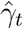 through the relation 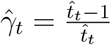 which relates the amount of past evidence seen to the weight that should be given to this past evidence. As expected given the monotonicity of this relation, the subjects’ approximate counts overestimate the true counts for *t <* 5, and underestimate them for *t >* 5 (Fig. 3, bottom right). As mentioned, this central tendency is a typical Bayesian pattern. But notably, we find that the shape of the approximate counts, as a function of the correct counts, resembles that obtained in numerosity studies, in which subjects are asked to estimate a number of items; in particular, the underestimation bias for the large counts is greater than the overestimation bias for the smaller counts ^22–24^. In short, it seems that the subjects in the averaging task make errors in their estimates of the numbers of observations in the same way that subjects in numerosity-estimation tasks make errors in their estimates of the numbers of items presented to them. As for the linear-bias model, it captures well these patterns (Fig. 3, green lines), although it seems to overestimate the bias of subjects for low counts (see Fig. 3, bottom left, for *t* = 2 and 3). This also points to the fact that subjects make smaller errors for low counts, presumably because after 2 or 3 observations, they still have a reasonably good idea of how many observations they have seen.

The four panels of Figure 3 show the same data under different lights: the successive weights 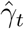 (top left), the bias 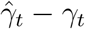 (bottom left), the number weights *w*_*t*_ (top right), and the approximate counts 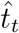 (bottom right) are all quantities directly related to each other (although the relation between 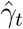 and *w*_*t*_ is a complex one; see Eq. 4). A common view is to consider the weights *w*_*t*_ (top right) and to point at two behavioral patterns: the primacy effect and the recency effect. We argue for a different view, represented in the other three panels. Under this view, the recall by subjects of the count of observations is imprecise, and on average, biased (in a way consistent with Bayesian inference); this leads them to weigh incorrectly, in their update of the average, their previous estimate and the new number. In turn, this results in numbers weights that exhibit both primacy and recency effects. These two effects are thus simultaneously accounted for by a single, plausible mechanism of imprecise recall.

One implication of this mechanism is that the crossover point *τ* should depend on the total number *T* of observations in a sequence. In our model we have chosen, for the crossover point, the midpoint between 2 and *T* = 8, i.e., 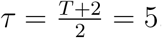, as a straightforward specification of a central tendency for count estimates. The effective crossover point in subjects’ responses seem indeed to be close to this value (Fig. 3). To test whether sequences of different lengths result in different crossover points, we turn to other datasets, that pertain to similar averaging tasks. The first dataset we consider comes from the same study as the one we have focused on so far^20^, but in this case the sequences are half as long, i.e., *T* = 4. We have collected the second dataset in the context of a different study ^25,26^, in which the sequences of number had length *T* = 5. For both studies we thus predict that the crossover point *τ* should be lower than the one posited so far, 5. We note, however, that the tasks in these two studies are different than the one we have considered so far: in both cases, subject were presented with two sequences of numbers (each of length *T*), and they were asked to compare the averages of the two (instead of observing one sequence and comparing its average to a fixed, reference value). Thus any effect we find may originate in this difference in task design. In fact, a possibility is that keeping track of two sequences comes with an increased cognitive load that may accentuate the imprecision in the recall of the counts, and this may lower the crossover point of the count estimates vs. the correct counts, thus introducing a potential confound for our predictions.

Notwithstanding this caveat, we compare the crossover point *τ* in these three datasets that differ by the sequence length *T*. Our prediction is that *τ* should be near the midpoint between 2 and *T*, and thus roughly obey the equality 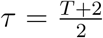. To estimate *τ* in each case, we fit the linear-bias model specified by Eq. 8 to the data, but this time we let *τ* be a free parameter. For the main dataset considered in this study (*T* = 8), the best-fitting crossover point is *τ* = 5.57, slightly higher than the value we have posited in our model. For the other two datasets, our predictions are *τ* = 3.5 for the dataset with *T* = 5, and *τ* = 3 for the dataset with *T* = 4. We find *τ* = 3.54 and *τ* = 3.16, respectively (Fig. 4). These values are reasonably close to our predictions. They do not significantly differ from each other, but they are both significantly lower than that obtained with *T* = 8. Across these three values of *T*, the crossover point *τ* thus seems to be an increasing function of *T*, as predicted.

**Fig. 4:**
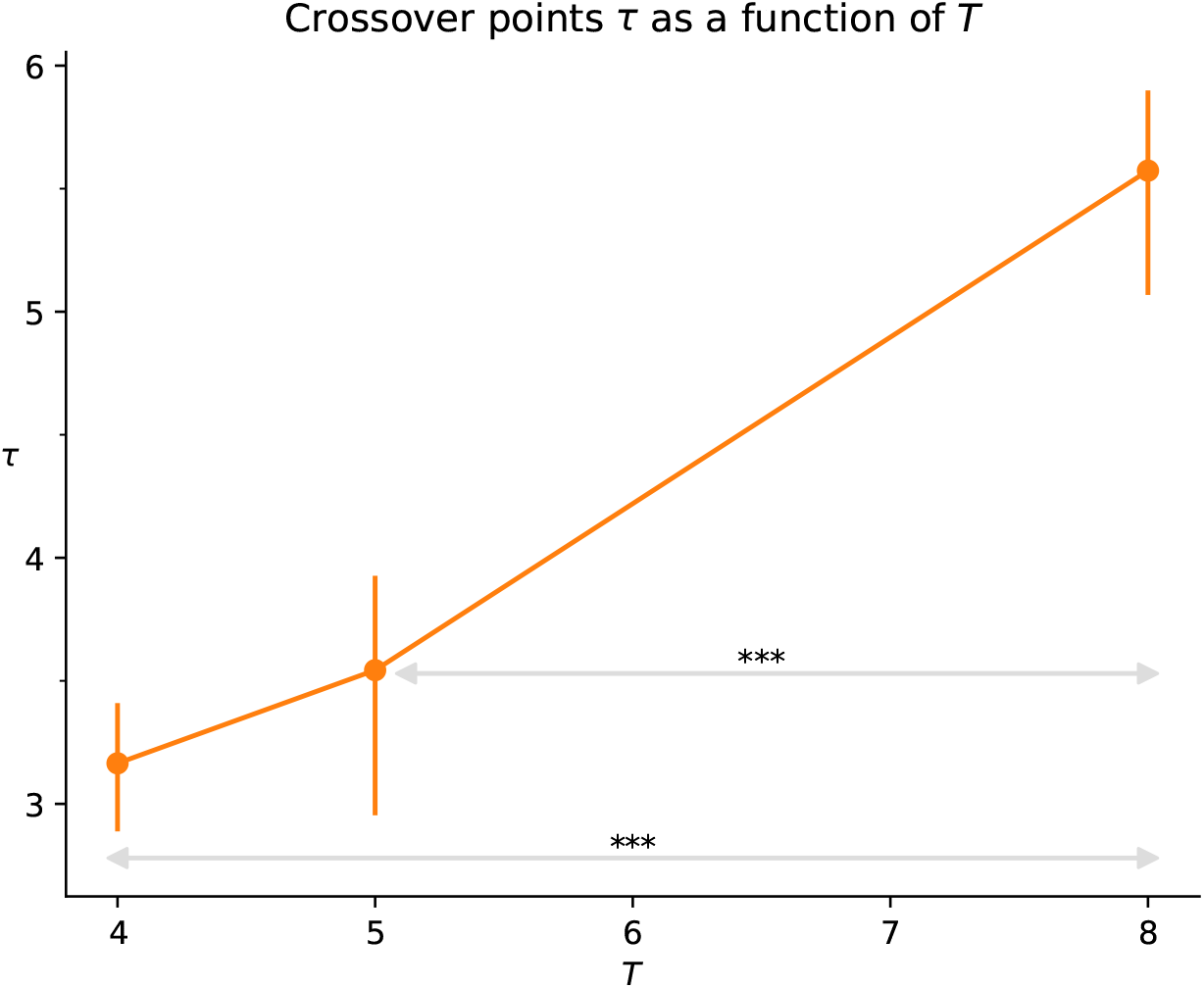
Subjects’ central tendency shifts towards lower counts when sequences are shorter. Subjects’ median crossover point *τ* as a function of sequence length *T* in different studies^20,25^. Stars indicate the p-values of permutation tests (30,000 permutations) of the equality between subjects’ crossover point for *T* = 8 vs. *T* = 4 and *T* = 5 (***: *p <* 0.001). Error bars indicate bootstrap-estimated 95% confidence interval.

## Discussion

We have introduced a model of imprecise recall of the count of seen observations that yields both primacy and recency effects in a sequential number-averaging task. Consistent with central tendency, a hallmark of Bayesian inference, subjects appear to overestimate the count of observations near the beginning of the sequence, leading to an overweighting of preceding numbers (primacy effect), and to underestimate the count toward the end of the sequence, giving too much influence to recent numbers (recency effect). This simple misevaluation of the count of observations provides a unified explanation for these seemingly distinct phenomena (Fig. 3). Our approach, moreover, nests exponential-discount models as a special case in which subjects have no sense of how many items they have seen.

Typical accounts of serial-position effects rely on a ‘leak’ term that yields an exponential discount which captures the ‘forgetfulness’ of the decision-making process^12,16,19,20^, and which can be seen as an optimal adaptation, either to the cognitive constraints hindering information integration^11,27^, or to the putative uncertainty in the environment^28,29^. This leak term usually results in recency effects, but it may yield primacy effects if allowed to be negative, which has been interpreted as a confirmation bias^13^. In memory recall experiments, in which subjects are asked to recall words that were presented to them sequentially, serial-position effects have been interpreted as reflecting the underpinnings of two storage mechanisms, one short-term and one long-term^4,6^. At first sight our account of serial-position effects, by which subjects do not know precisely how much weight they should apply to a new number in computing an updated average, may seem to be only relevant for the specific task of averaging numbers, with little if any implications for other domains. We note however that in a sequential inference task, we have observed that the way people update their beliefs was precisely largely insensitive to the amount of evidence previously seen, suggesting that there also subjects did not correctly weigh past and new information ^30^.

Thus we argue that the mechanism we introduce applies more generally than solely to averaging tasks. To further illustrate this point in the context of memory-recall tasks, we sketch a model that is in spirit very similar to the model we have presented. Suppose that an observer is presented with *T* words, sequentially, and has at their disposal *N* bits to store these words. The more bits dedicated to store a word, the more likely is the observer to recall accurately this word. We denote by *n*_*i,t*_ the number of bits dedicated at time *t* to the word that was presented in position *i* ≤ *t*. We also posit that the observer always uses all the bits available, i.e., 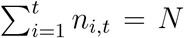, for all *t* ≤ *T*. Thus upon receiving the word in position *t >* 1, all bits are ‘taken’, but our observer decides to dedicate a fraction 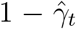 to the new word, i.e., 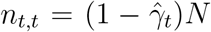, and to trim the memory of all the previous words by the same factor, i.e., 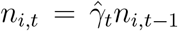 (thus still using all the bits). After receiving the last word, the fraction of bits dedicated to each word depends on the sequence of chosen fractions, 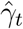, and is equal to the weight *w*_*t*_ given in Eq. 4, i.e., *n*_*i,T*_ = *w*_*i*_*N*. If the observer chooses 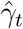 as equal to 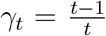, then the *N* bits are at all time equally divided among the *t* words seen; but this requires remembering the count *t*. If the observer has only access to an approximate estimate of this count, then the fraction *w*_*t*_ dedicated to each word, as a function of *t*, will exhibit the non-monotonic, U-shape behavior shown in Fig. 2F — and so will the probability of recalling each word. Thus this observer exhibits primacy and recency effects in a memory-recall task.

We leave the study of this memory model for future work. We surmise, moreover, that a similar approach could be applied to other domains. Finally, we note that our model predicts that two factors should influence the primacy and recency effects in sequential tasks. First, the distribution of the total number of stimuli *T* in a sequence (in the datasets we have considered, *T* was fixed, and we have provided some evidence that subjects are indeed sensitive to it; see Fig. 4). Second, the precision with which subjects estimate the count *t*, which could be manipulated by changing the duration of presentation, or by increasing the cognitive load of the subject (e.g., by asking the subject to complete another task in parallel). We leave these further investigations of the model to future studies.

## Notes

### Competing Interest Statement

The authors have declared no competing interest.

